# Establishing neuroanatomical correspondences across mouse and marmoset brain structures

**DOI:** 10.1101/2024.05.06.592808

**Authors:** Christopher Mezias, Bingxing Huo, Mihail Bota, Jaikishan Jayakumar, Partha P. Mitra

**Affiliations:** Cold Spring Harbor Laboratory, 1 Bungtown Rd, Cold Spring Harbor, NY; Broad Institute of MIT and Harvard, Data Sciences Platform Division, 105 Broadway, Cambridge, MA; Indian Institute of Technology-Madras, Center for Computational Brain Research, Chennai, TN, India

## Abstract

Interest in the common marmoset is growing due to evolutionarily proximity to humans compared to laboratory mice, necessitating a comparison of mouse and marmoset brain architectures, including connectivity and cell type distributions. Creating an actionable comparative platform is challenging since these brains have distinct spatial organizations and expert neuroanatomists disagree. We propose a general theoretical framework to relate named atlas compartments across taxa and use it to establish a detailed correspondence between marmoset and mice brains. Contrary to conventional wisdom that brain structures may be easier to relate at higher levels of the atlas hierarchy, we find that finer parcellations at the leaf levels offer greater reconcilability despite naming discrepancies. Utilizing existing atlases and associated literature, we created a list of leaf-level structures for both species and establish five types of correspondence between them. One-to-one relations were found between 43% of the structures in mouse and 47% in marmoset, whereas 25% of mouse and 10% of marmoset structures were not relatable. The remaining structures show a set of more complex mappings which we quantify. Implementing this correspondence with volumetric atlases of the two species, we make available a computational tool for querying and visualizing relationships between the corresponding brains. Our findings provide a foundation for computational comparative analyses of mesoscale connectivity and cell type distributions in the laboratory mouse and the common marmoset.

## Introduction

The mouse (*Mus musculus*) is the most widely used mammalian species in neuroscience research^1,2^. Studies have compared mouse and human brains in terms of anatomy^3^, functional connectivity^4^ and gene expression^5^. However, instead of providing a conclusive comparative framework, these studies have highlighted the major challenges in translating experimental results from mice to humans^6,7^. As a result, there has been a growing trend in research programs that use alternative models^8^, with a particular focus on non-human primates^9^, increasingly the common marmoset (*Callithrix jacchus*)^10^. Non-human primates have long been recognized as important in neuroscience research because of their physiological similarities and evolutionary proximities with humans^11^. While traditionally studied for aspects of their social behaviors, such as vocalization, common marmosets have attracted increased interest as animal models due to the relatively short time taken to reach reproductive maturity large litter sizes compared to macaque monkeys, and the development of molecular genetic tools^12,13^. Thus, marmosets form a natural bridge between mouse models and humans for basic and translational research^14,15^. Furthermore, modern neuroscience techniques first developed in mice are increasingly being applied to marmosets, opening the possibility of substantive cross-species comparisons^16,17^.

However, before conducting refined comparative analyses on brain connectivity or cell-type distributions beyond major brain compartments, a crucial issue must be addressed: establishing correspondences between the brains of the two species. To some extent, this is a chicken and egg problem, since the connectivity and cell-type distribution data is needed to pin down such correspondences. Nevertheless, there is a need for a starting point for recursive refinement, based upon the literature and neuroanatomical knowledge. This is the problem we address in the current work. We propose a conceptual framework to establish correspondences between reference atlases for mouse and marmoset and use this framework to generate a concrete mapping between named brain compartments from the two species. This is reminiscent of the alignment of genomes of different species, but substantially more challenging. Unlike genomes, fragments of which are sufficiently conserved across species to permit a base-pair level alignment in places, precise spatial alignment at the microscopic level of individual neurons is not possible even within individuals of a given species, let alone between animal taxa. Such alignment or mapping between brains of different taxa is only possible at the mesoscopic scale^18^, which roughly corresponds to named regions in histological atlases. We therefore focus on relating histological atlases across the two taxa at the level of named brain compartments.

The traditional method for identifying specific brain regions involves consulting corresponding histological atlases, which delineate the origins and targets of axonal tracts or quantify the spatial distribution of specific cell types. These atlases, typically in the form of printed books, overlay histological images with stereotactic coordinate grids, segmented into named regions. Computationally accessible three-dimensional reference atlases that organize brain regions in a nested hierarchy^19–21^ are available for some species. Ideally, different brain atlases within each species should be in good agreement, so that one can focus on the cross-species mapping. Unfortunately, this is not the case. Even within a species, diverse atlases have emerged to address specific research questions or techniques, resulting in heterogeneous segmentations of the same brain regions and distinct hierarchical organizations of named compartments without much standardization of names. Efforts have been made, e.g. by Paxinos and colleagues, to harmonize named compartments across mammalian brains^22–25^, but other atlases remain prevalent, and the problem is far from resolved. It is worth noting that atlases evolve with our knowledge and technology, contributing to inconsistencies in segmentation and organization of the same brain regions in the same species across different versions and atlases.

Establishing correspondences between brain atlases of mouse and marmoset is therefore a complex, dynamic and challenging process. Within-species mappings between brains of different individuals, using diverse imaging modalities, resolutions and anatomical variations, is usually handled using multimodal brain data registration algorithms^26^. Within-species volumetric atlases may be reconciled algorithmically using the volumetric overlap between different brain compartments^27^. For cross-species comparisons, this is not a workable strategy since the reference brains have quite different geometries, and it is unrealistic to expect that a smooth spatial mapping can be developed between mouse and marmoset brains that is valid at the smaller compartment scales. Such an approach might become more feasible if a much larger set of brain atlases of many taxonomic groups and species becomes available in the future, so that the mapping can be done pairwise through “nearby” brains on the phylogenetic tree. But this is not feasible today. Approaches based on literature mining have been proposed^28^. Homological relations based on common ancestry cannot be easily inferred from any single observable characteristic^29,30^, so methods for identifying homologues in comparative neuroanatomy are complex^29,31–33^. Several databases based on literature studies have clustered nomenclature of potentially homologous regions in different species and organized them into hierarchical structures. Examples include NeuroNames^34–36^, Brain Architecture Management System (BAMS)^37,38^, Neuroscience Information Framework (NIF)^39,40^ and Uber-anatomy ontology^41,42^. The heterogeneity across these resources poses challenges to the investigator wishing to compare brain connectivity architecture or cell type distributions in the extensive data sets that are now available and emerging for the mouse and the marmoset.

Our proposed approach is based on our empirical observation that disagreements among neuroanatomists about establishing correspondences between brain parts primarily stem from differing opinions on the hierarchical, tree-like organization that should be applied to the smallest compartments. Traditional wisdom holds that higher level brain compartments should be easier to homologize since they are evolutionarily older structures. To some extent, this is true; specifically, there is little debate about establishing correspondences between, for example, the cerebelli of different taxa. However, establishing correspondences for such macroscopic compartments does not provide scientific insights, as these compartments are typically highly spatially heterogeneous. By examining the literature and existing atlases, we discovered, somewhat paradoxically, that the opposite of the conventional wisdom is true. We found that plausible correspondences between mouse and marmoset brains were easier to establish at the “leaf-levels” of the hierarchically structured parcellations of these brains, corresponding to the smaller structures. For example, there is little disagreement in what the amygdala is in the mouse and the marmoset; however, there is significant disagreement in where to put the amygdala in the next level of the hierarchical tree. Our solution is therefore to discard the hierarchy and confine our attention to the leaf-level structures to establish the correspondence between the two species’ brains.

We identified correspondences between the most granular level, or leaf-level, structures in atlas hierarchies of the mouse and the marmoset, based on different existing atlases, databases and literature for these species. The atlases we utilized include the most used atlases for mouse (Allen Mouse Brain Atlas^43^ and Paxinos Mouse Brain Atlas; see **Extended data Table 2**; rows 1,2), and marmoset (Paxinos Marmoset Brain Atlas^24^ and the RIKEN Brain Atlas; see **Extended data Table 2**, rows 3,4). We first interpolated between atlases within the same species to get a single unified set of leaf-level regions for both mouse and marmoset (Supplementary materials) and then assigned correspondences between these leaf-level structures based on an examination of the atlases and the literature. Our primary resulting output is a full list of corresponding leaf-level brain regions between the two species (Supplementary materials), thus enabling the establishment of comprehensive interspecies homologies across the entire brain. This is a challenging endeavor, involving the manual examination of thousands of named compartments across the two species in four atlases and over one hundred original publications. We categorise and analyze the types of correspondences (1 to 1, 1 to many, many to 1, and many-to-many, as discussed below in the results section and Table 2). We were able to place 268 of the 627 named mouse leaf level structures (43%) in 1-1 correspondence with a corresponding 268 of the 569 marmoset leaf level structures (47%). Of the remainder, 45 mouse structures (7%) had a one-to-many correspondence with 135 marmoset structures (24%), potentially indicating differentiation in the marmoset since many of these 1-many correspondences were of cortical compartments. Vice versa, 104 (17%) of mouse structures were in many-1 correspondence with 33 (6%) marmoset structures. 25% of the structures in mouse and 10% of those in marmoset did not have evident counterparts in the other species, whereas the remaining structures were involved in small groups of many-to-many relationships.

We expanded this analysis to volumetric atlases containing segmented volumes, using the Allen atlas (**Extended data Table 2**; row 1) for mouse, and a refined version of the RIKEN atlas (**Extended data Table 2**, row 4) for marmoset. Finally, we develop a MATLAB package, downloadable from GitHub, which allows users to query, via a GUI, the correspondences between the two species’ volumetric atlases. We expect that the comparative framework established here, along with the MATLAB package, will facilitate comparative neuroanatomy between the mouse and marmoset, at a whole-brain scale, regardless of experimental technique or data modality. It will allow us to understand what aspects of the connectivity architecture or cell type distributions are common across the taxa, and what aspects reflect taxonomic differences possibly related to niche differentiation.

## Results & Discussion

### Cross-species atlas reconciliation via leaf-level brain compartment homological correspondences

Two mouse brain atlases and two marmoset brain atlases were used in the current study: the Allen Mouse Brain Atlas^44^, referred to as “Allen atlas” (**Extended data Table 2**, row 1); the Paxinos Mouse Brain Atlas (**Extended data Table 2**, row 2), referred to as “Paxinos mouse”; the Paxinos Marmoset Brain Atlas (**Extended data Table 2**, row 3), referred to as “Paxinos marmoset”; and the Marmoset Brain Atlas from the Brain/MINDS project^45^, referred to as “RIKEN atlas”. Because previous methods developed for within-species atlas comparison lack cross-species translational power, we approached reconciliation of the different species’ atlases by identifying correspondences between leaf-level brain compartments (**Table 1; Supplementary Table 1**). Of note, the RIKEN atlas nomenclature is a subset of the Paxinos marmoset nomenclature, so the list of brain regions considered for comparison is one rather than two columns. The RIKEN atlas is necessary to include as it provides a volumetric segmentation of marmoset brain regions.

**Table 1.**
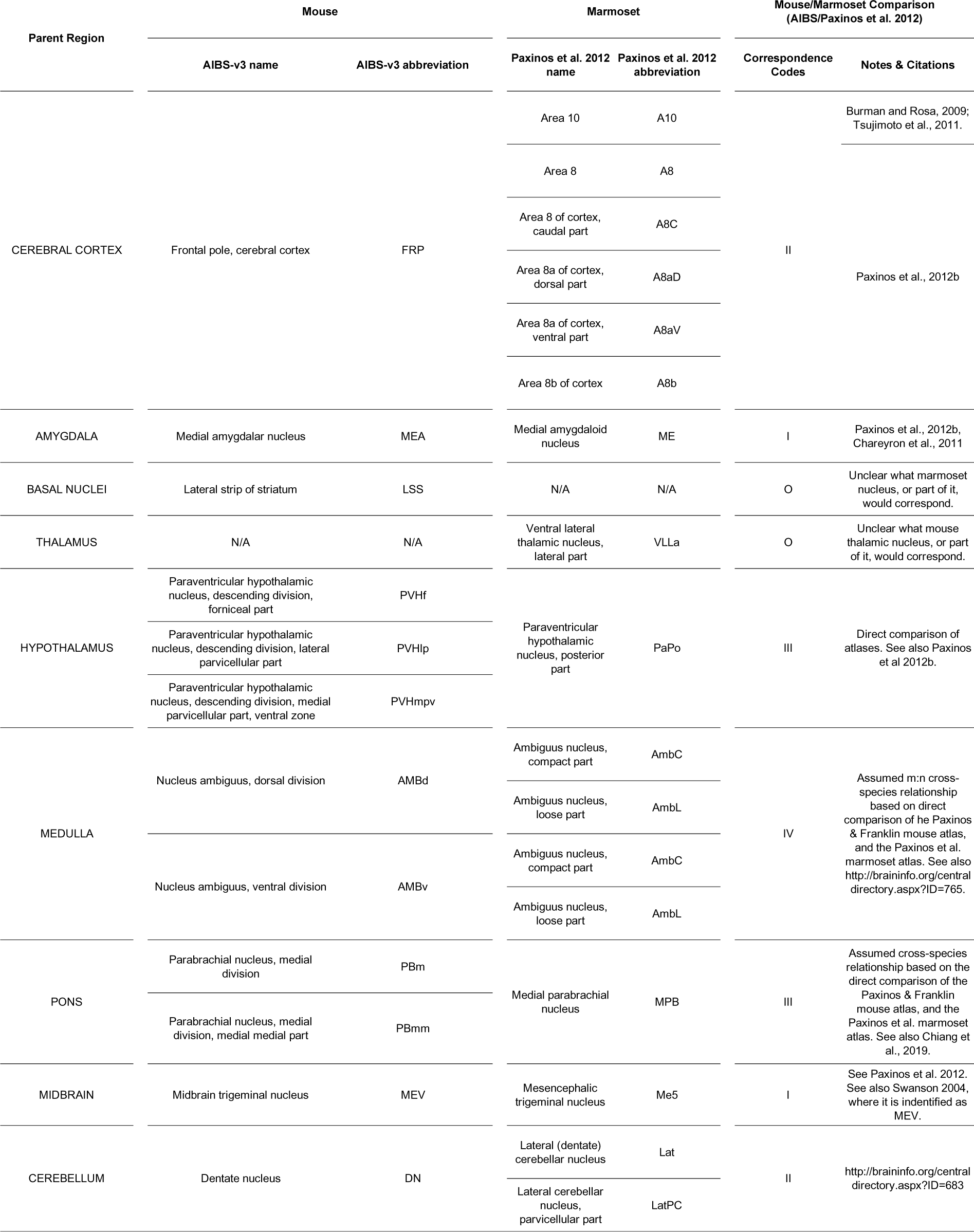
Excerpts of the full Supplementary Table 1 showing the different types of correspondences between mouse and marmoset nomenclatures. Note that this only shows a subset of hand-selected brain regions and only Allen Mouse and Paxinos marmoset correspondences. The fuller **Supplementary Table 1** contains correspondences between all mouse and marmoset atlases employed, and **Supplementary Tables 2, 3** list the leaf-level brain regions employed in the two species.

Our first task was to establish a flat list of leaf-level structures within each species corresponding to a partitioning of the brain (**Supplementary Table 2**). We primarily defined the leaf-level structures as finest granularity of brain compartments in a given atlas or nomenclature, with some caveats. First, we did not subdivide cortical layers; once a regional correspondence can be established, further division into cortical layers is a less challenging task. On occasion, we adopted as the leaf-level structure, the parent structure that is one level above the most granular parcellation, if this significantly facilitated making correspondences. When determining the set of leaf-level structures for mouse brain regions, special care had to be taken as we had to interpolate between both Allen and Paxinos mouse atlases. We considered 3 cases: 1) the Allen atlas contained only a parent region from the Paxinos mouse atlas, in which case the Paxinos mouse atlas regions were considered leaf-level; 2) the Paxinos mouse atlas contained only a parent region from the Allen atlas, in which case the Allen atlas regions were considered leaf-level; and 3) both contained partially or completely non-overlapping parcellations at a leaf-level, in which case we gave the Allen atlas regions primacy, as this atlas contains a volumetric segmentation. Note that we also provide a within-mouse comparison between Allen and Paxinos mouse atlases in **Table 1** and **Supplementary Table 1**.

The bulk of our effort consisted of manually establishing cross-species correspondences between the leaf-level structures we established (**Supplementary Tables 2, 3**). We inferred these correspondences from previous studies and existing databases^32^ as well as an examination of the corresponding histological atlas plates. Six types of correspondence were identified. Type O: no corresponding regions found in the other species. Type I: one-to-one corresponding regions between species. Type II (one to many): one leaf-level region in the mouse corresponds to multiple regions in the leaf-level marmoset list, due to a finer parcellation in the marmoset, which could potentially indicate specialization and differentiation in the marmoset. Type III (many to one): the reverse situation, where one marmoset leaf-level structure corresponded to multiple mouse structures. Type IV (many to many): multiple regions in the combined mouse ontology correspond to multiple regions in the marmoset ontology, possibly due to different parcellation schemes applied in the different species. An additional flag, Type U, indicates that correspondence between a given set of mouse and marmoset regions potentially exist, but the precise correspondence is uncertain. **Supplementary Table 1** shows the full list of all leaf-level structures and their correspondences in the two atlases, with an example subset given in **Table 1**.

From our leaf-level brain region correspondence analysis in mouse and marmoset, 466 out of 627 mouse leaf-level gray matter regions found correspondences, in one of the above categories, to 510 out of 569 marmoset leaf-level gray matter regions (**Table 2**). Approximately 43% and 47% were of Type I concordance between mouse and marmoset, indicating a large proportion of 1:1 relationships between the two species’ ontologies. These 1-1 relations are dominant in thalamic, midbrain and hindbrain structures but less so in cortical regions, which makes sense given that greater evolutionary divergence is expected at the level of cortical regions. Type II concordance, where multiple marmoset leaf-level regions map to one mouse leaf-level region, were the determined relationship in about 7% of mouse regions and 24% of marmoset regions, driven largely by cortical areas as the marmoset cerebral cortex is significantly expanded and differentiated compared to the mouse brain. 17% of mouse regions and 6% of marmoset regions were of Type III concordance, indicating finer parcellation among the corresponding compartments in the mouse as compared with the marmoset. This could either be due to rodent-specific specializations in the mouse brain, or due to a closer study of the mouse brain in past research studies leading to these refined parcels. Type IV correspondence accounted for about 4% of mouse regions and 9% of marmoset regions and was found in the amygdala, and in some cortical and thalamic areas where homology at the finest level was more challenging to establish.

**Table 2.** Breakdown of leaf-level brain region correspondences between mouse and marmoset atlases ontologies, using the Allen mouse, Paxinos mouse, Paxinos marmoset, and RIKEN marmoset atlases (**Extended data Table 2**, rows 1-4). Codes indicate relations from mouse to marmoset and are defined as: I (1:1); II (1:n from mouse to marmoset); III (n:1 from mouse to marmoset); IV (m:n from mouse to marmoset); O (no correspondence); I* (1:1, but uncertain); IV* (m:n, but uncertain).

| Leaf-Level Correspondence | Mouse |  | Marmoset |  |
| --- | --- | --- | --- | --- |
|  | Count | Percent | Count | Percent |
| **Total** | 627 | 100.00% | 569 | 100.00% |
| **O** | 161 | 25.68% | 59 | 10.37% |
| **I** | 268 | 42.74% | 268 | 47.10% |
| **II** | 45 | 7.18% | 135 | 23.73% |
| **III** | 104 | 16.59% | 33 | 5.80% |
| **IV** | 28 | 4.47% | 52 | 9.14% |
| **I\*** | 12 | 1.91% | 12 | 2.11% |
| **IV\*** | 9 | 1.44% | 10 | 1.76% |

Type O correspondence reflects the lack of evidence for potentially homologous areas between the two species. While this is true for most Type O areas in our results, we note the following caveats. First, although the correspondence was not identifiable between the mouse and marmoset atlases, it could be identifiable in other atlases. For example, the commissural nucleus of the inferior colliculus in marmoset did not find a corresponding region in the mouse atlases. Yet, a rodent counterpart can be identified in the rat atlas^46^ as the commissural nucleus of the PAG (peri-aqueductal grey), implying its existence in mouse. Second, corresponding areas were found in the literature but not present in the atlases. Examples include the retroreuniens nucleus in marmoset, whose mouse counterpart was not found in the mouse atlases, but other literature indicated overlap with the caudal part of the medial reuniens nucleus. Similarly, retroisthmic nucleus in the marmoset does not have a clear mouse counterpart in the utilized mouse atlases, but the term was found in the literature^47^. Conversely, submedial nucleus of the thalamus (SMT) in mouse has its counterpart in primate species according to previous work^48,49^ despite a lack of annotation in the employed marmoset atlases. We performed a thorough examination of these cases and included them in **Supplementary Table 1**.

### Potential anatomic and neuroscientific implications of leaf-level brain homological concordances

Correspondences were drawn from the existing literature and databases to establish reasonable comparisons and formulate testable hypotheses. For example, subdivisions of retrosplenial cortex in mouse vary across atlases^50,51^, leaving unclear correspondences between these substructures in the Allen atlas and subdivisions of Areas 29 and 30 in the RIKEN atlas. We moved one level up and drew correspondences between the retrosplenial cortex in mouse and Areas 29 and 30 in marmoset. One hypothesis associated with this correspondence is that the corresponding regions in the two species are homologous even if their substructures are non-homologous, following the argument in Striedter & Northcutt^31^. As another example, both temporal and parietal association cortices are involved in higher-order processing of perceptual, cognitive and motor functions in anthropoids^52^. Compared with a single region of temporal association area in the Allen mouse atlas, the marmoset temporal association area contains 7 substructures, in addition to a “lateral and inferior temporal cortical region,” which is further divided into substructures. While the posterior parietal association area is considered an integral area together with anterior and rostrolateral visual area in the mouse atlas, it is subdivided into 10 substructures in the marmoset atlas. One hypothesis associated with this correspondence is that the temporal and parietal association cortices in both species evolved from the same ancestral brain region, and the higher complexity and fractional expansion in the marmoset brain is related to certain gains-of-function in marmosets or losses-of-function in mice^52^.

### Leaf-level regions provide a more stable basis for cross-species comparative anatomical and ontological analyses than larger parent region groups

We demonstrate the idea that leaf-level correspondences are better suited for interspecies comparisons. The hierarchies of gray matter regions in the two volumetric brain atlases (Allen mouse and RIKEN marmoset) form tree structures, where the whole brain gray matter is the tree root, and the most granular brain compartments are leaves of the tree. First, we show that the tree structures are very different for the two atlases, implying that the grouping of leaf-level regions into parent regions differ between the two. This reflects distinct criteria of organization used by the expert neuroanatomists (**Fig. 1A, C**). Criteria used to define the hierarchical structure of the Allen/Swanson atlas (**Fig. 1B**) include gene expression clustering^44^ and adult brain function^50,53^, whereas the construction of the RIKEN/Paxinos atlas hierarchy (**Fig. 1D**) gives increased weight to mammalian brain developmental processes^54,55^. There is no clear-cut way to choose between these organizations, since any atlas would need to be draw upon much previous work, covering many different fields of neuroscience. Different organizing principles in creating these hierarchies and in defining parent regions pose challenges to studies leveraging either species, hindering cross-species comparison.

**Fig. 1.**
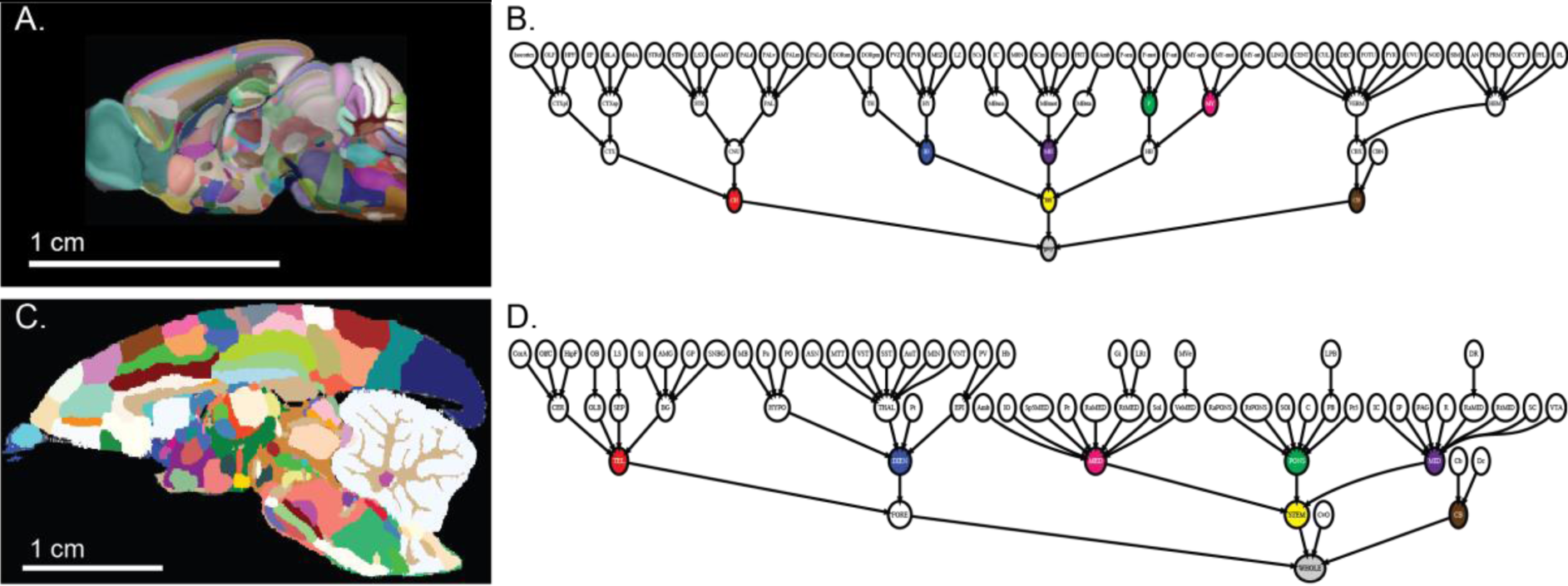
**A**. A sagittal view of the Allen mouse reference brain with CCF3 segmentation. **B**. High-level branches of the Allen atlas hierarchy. **C**. A sagittal view of the Brain/MINDS marmoset reference brain with refined segmentation. **D**. High-level branches of the marmoset brain atlas hierarchy. Color coded nodes in the trees: red: cerebrum/telencephalon; blue: interbrain/diencephalon; yellow: brain stem; green: pons; purple: midbrain; magenta: medulla oblongata; brown: cerebellum.

One of our primary observations is that the hierarchical grouping of the leaf-level structures may be a distracting factor when establishing correspondences. In fact, once a leaf-level correspondence has been established, any hierarchical tree structure may be superposed on top, without removing the ability to do comparative analyses at the granular level. To demonstrate this, we took a set of leaf-level regions with Type I-IV correspondence and superimposed different hierarchical organizations of brain regions based on different criteria used by different neuroanatomical experts (Paxinos^55^, Swanson^50^). **Fig. 2** shows that different hierarchical trees from the Allen and RIKEN atlases may be superimposed onto a set of corresponding leaf-level regions. We could consequently derive different sets of parent-level regions, corresponding to different organizing principles and metrics, while still retaining the same set of leaf-level regions as the basis for cross-species comparisons. Therefore, reframing cross-species anatomical comparisons around leaf-level regions avoids potentially controversial decisions about organizing principles when defining parent regions and hierarchies, while allowing for meaningful analysis. If comparisons need to be drawn for larger structures, such superstructures may be chosen that correspond to the same set of leaf level structures, for connectivity of cell-type distribution studies.

**Fig. 2.**
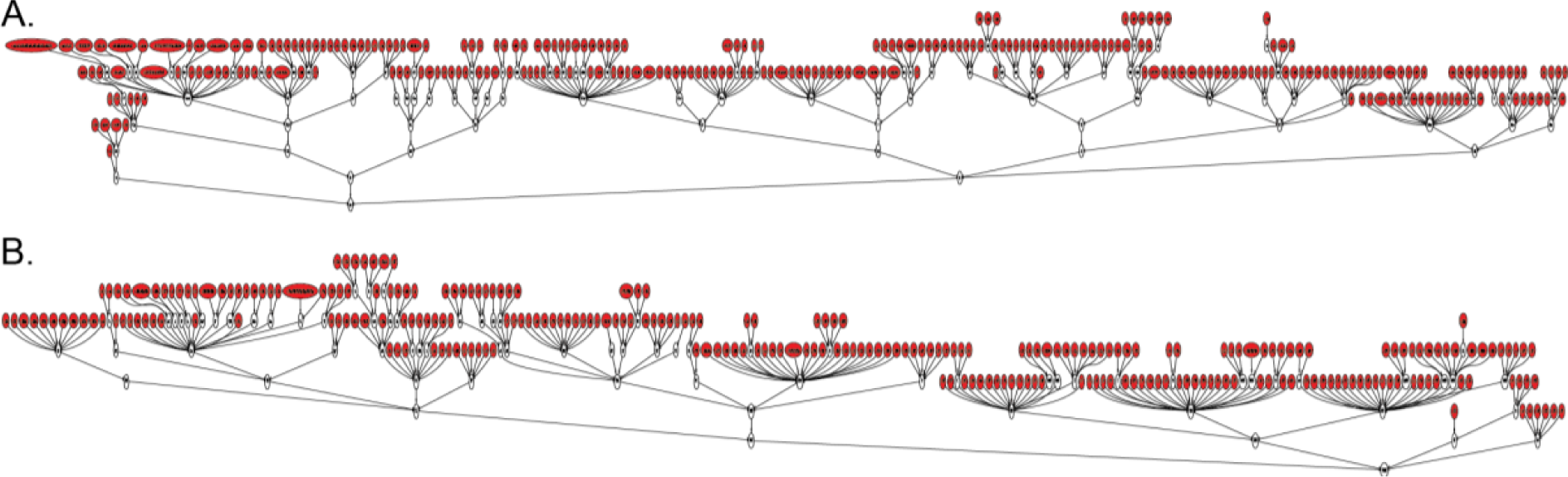
**A**. The Allen atlas hierarchy superimposed on marmoset leaf-level regions (red). **B**. The RIKEN atlas hierarchy superimposed on mouse leaf-level regions (red).

### Cross-species correspondence of segmented volumetric atlases

The Allen atlas (**Fig. 1A**) and RIKEN atlas (**Fig. 1C**) provide segmented volumes matching their hierarchically structured nomenclatures (**Fig. 1B, D**), allowing spatial, in addition to purely nomenclature-based, analyses. Leveraging the cross-species correspondences and segmented brain volumes, we compared volumes of brain regions with homologous counterparts. The Allen mouse reference brain had whole-brain gray matter segmentation (CCF3)^56^ based on the Allen atlas nomenclature and hierarchy. However, the marmoset reference brain accompanying the RIKEN atlas only had 63.5% of the total volume annotated^57^. To mitigate this drawback, we complemented the Brain/MINDS marmoset reference brain segmentation with NIH-Silva and Saleem marmoset reference brain segmentations^58^ (**Extended data Table 2**, rows 5, 6), by co-registering reference brains from these segmentations to a common reference space to which we had mapped the RIKEN segmentation. This common reference space was derived from the average of 43 T2-weighted MRI scans of female marmoset brains (**Fig. 1C**).

Fractional volumes of common key regions in the Allen and NIH-Silva Marmoset and Saleem Marmoset refined RIKEN atlases can be evaluated by combining the reconciled atlas leaf-level regions and the 3D reference brains (**Table 3)**. 98.78% of Allen labeled volume corresponded to gray matter; however, this number was only 78.3% in marmosets. This could be either an effect of differences in regional assignments between atlases or a real biological difference, as primate brains generally have dense gray matter paired with large white-matter tracts. Leaf-level regions accounted for 82.5% and 87.1% of labeled mouse and marmoset gray matter, respectively. The remaining ∼15% of gray matter in each atlas consists of non-specific labels, e.g. “Medulla” or “Cerebellum”; such labels were not compared to establish correspondences but do count towards the total gray matter volume. Akin to the results in **Table 2** based on comparing compartment names, we find (**Table 3**) that Type I relations account for the largest fraction of gray matter volume of both species, but not to the same degree. Interestingly, Type II (1 mouse:n marmoset) relations account for a larger fraction of regions and gray matter volume in marmosets than Type III (n mouse:1 marmoset) relations in mouse. This suggests homological marmoset brain regions are more likely to be subdivisions of a larger region in mouse; much of this subdivision in marmoset occurs in cortical regions. Type IV and IV* relations (many:many) also account for a larger fraction of marmoset than mouse gray matter volume (**Table 3**); this difference is again driven by cortical regions. Taken together, the prevalence of Type II, Type IV, and Type IV* relations in marmosets suggests increased subdivision in cortical regions in marmoset compared with mouse.

**Table 3.** Comparison of mouse and marmoset volumetric atlases, using the Allen Mouse and RIKEN Marmoset volumes (with refinement from the NIH-Silva Marmoset and Saleem Marmoset segmented volumes). Total “Pct. Volume” is the percentage of the volume with gray matter labels, excluding ventricle and white matter labels. Subsequent rows of “Pct. Volume” give the percent of labeled gray matter volume per correspondence.

| Leaf-Level Correspondence | Mouse |  |  | Marmoset |  |  |
| --- | --- | --- | --- | --- | --- | --- |
|  | Count | Pct. Regions | Pct. Volume | Count | Pct. Regions | Pct. Volume |
| **Total** | 337 | 100.00% | 98.78% | 319 | 100.00% | 78.33% |
| **O** | 121 | 35.91% | 10.67% | 64 | 20.06% | 12.28% |
| **I** | 132 | 39.17% | 33.28% | 132 | 41.38% | 26.96% |
| **II** | 22 | 6.53% | 18.47% | 69 | 21.63% | 21.53% |
| **III** | 35 | 10.39% | 11.70% | 17 | 5.33% | 6.11% |
| **IV** | 22 | 6.53% | 5.38% | 34 | 10.66% | 17.38% |
| **IV\*** | 5 | 1.48% | 3.01% | 3 | 0.94% | 2.79% |

### Volumetric comparisons of cortical regions

In both Allen mouse and RIKEN marmoset reference brains, the cerebral cortex was fully segmented at the leaf-level. All of the cortical volume in mouse and 83.8% in marmoset found correspondences in the counterpart species. The marmoset cortical compartments that did not find correspondences in the mouse brain partially reflected the lack of homology of some cortical areas, and the different regional assignments such as piriform and entorhinal cortices, as mentioned above. We defined the cortical volumed as collections of key regions (with Type I-IV correspondences) that were assigned as cortex in both Allen and RIKEN atlases, as well as all Type O leaf-level regions in each species that were assigned to cortex. In total, the cortical areas comprise 123.3 mm^3^ or 27.4% of the total gray matter in the mouse brain, and 3288.4 mm^3^ or 57.2% of the total gray matter in the marmoset brain. By comparing the fractional volume of cortical areas in the two species (**Fig. 3**), we observed the most pronounced differences in the visual cortex, comprising 38.1% of the marmoset cortex, but only 10.9% in mouse. Similarly, the parietal and temporal association cortices occupy 7.1% and 14% of marmoset cortex, respectively; but only 2% and 2.5% in the mouse brain cortex. On the contrary, the motor and somatosensory cortices comprise 19.8% and 27.0% of the mouse brain cortex, respectively; but only 5.7% and 6.1% in marmoset. The retrosplenial cortex occupies 8.5% of the mouse cortex and 1.6% in the marmoset cortex. The insular cortex, including agranular and granular cortices, along with other regions that are marmoset-specific, occupies 8.3% of the mouse brain cortex and 2.1% of the marmoset brain cortex.

**Fig. 3.**
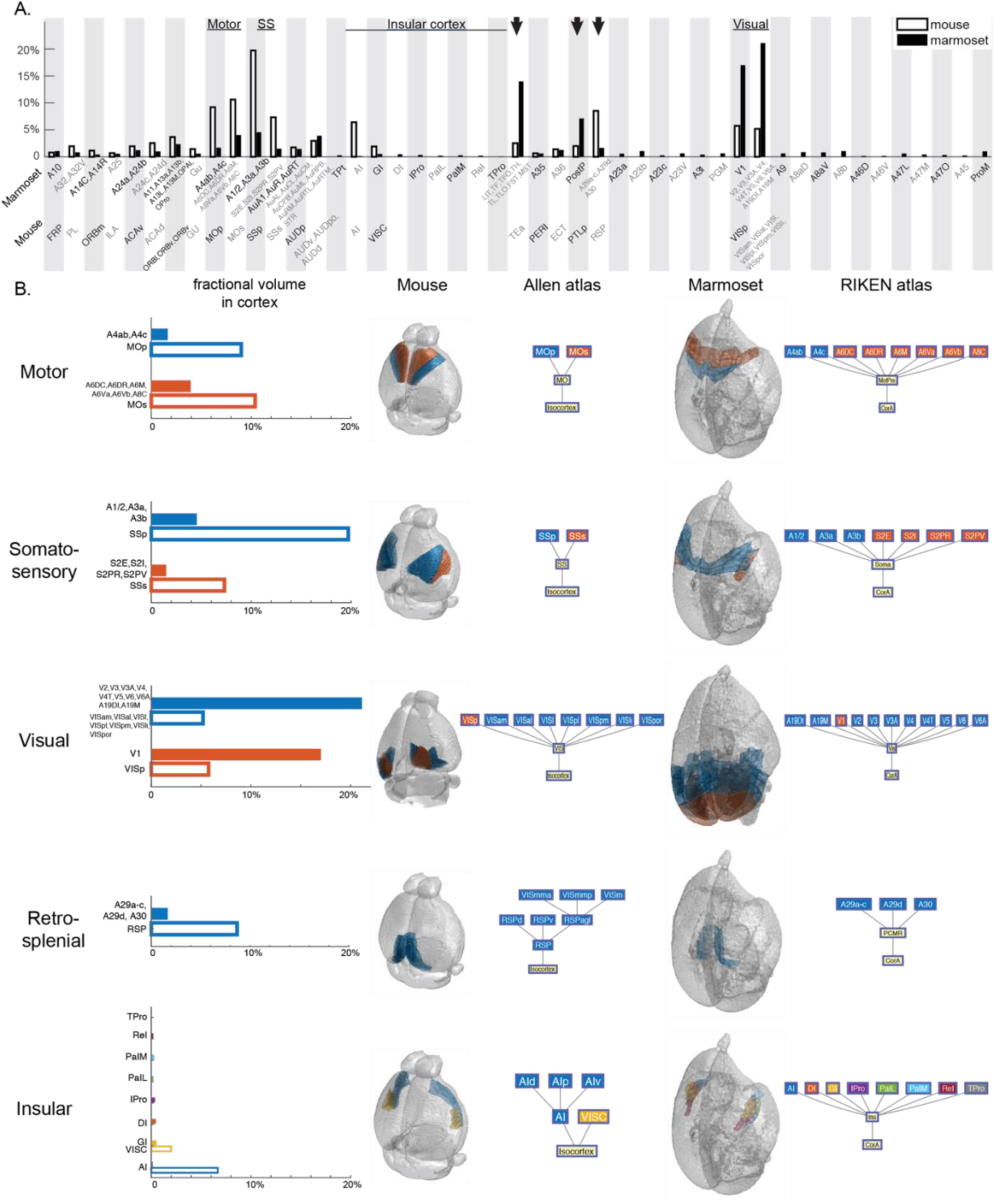
**A**. Fractional volume of some leaf-level cortical structures in the mouse (not filled bars) and marmoset (filled bars) cortices. Arrows point to retrosplenial, temporal and parietal association cortices. **B**. Sample cortical regions’ fractional volume in cortex in mouse and marmoset (left), in mouse reference brain and Allen atlas hierarchy (middle), in marmoset reference brain and RIKEN atlas hierarchy (right). For a full list of abbreviations and brain regions in both species, refer to **Supplementary Table 1**.

Within the redefined cortical volume, 11.5% of the volume or 26 cortical regions in the RIKEN atlas did not have correspondences in the Allen atlas (**Fig. 3**). These regions are mostly in the insular cortex (1.6% of the cortex volume), precentral opercular cortex (1.0%), Brodmann areas 8 (2.7%) and 23 (2.5%), and other smaller regions. These marmoset-unique regions had homologies in macaque brain atlases^54^, suggesting primate-rodent taxonomic differences. Note that navicular nucleus of the basal forebrain and supracallosal subiculum are leaf-level structures in the marmoset cortex but were excluded from the volume analysis since they were not annotated in the reference brain.

### Anatomic and neuroscientific implications of volumetric comparisons between mouse and marmoset brain atlases

Our quantitative evaluation of brain region volumes between species produced results that potentially reflect the evolution of functional specialization^59,60^. For example, we observed the visual areas occupying a larger volume fraction in marmoset cortex than in mouse cortex, and the reverse was true for motor and somatosensory areas. These observations are consistent with the living habitat and survival niche of marmosets, which are diurnal and arboreal, relying on vision for spotting food and prey, as well as mice, which are nocturnal and terrestrial, relying on tactile information to explore environments^59,61^. The insular cortex is involved in a wide range of functions including visceral, somatosensory, motor and limbic integration^62–64^, and occupies a larger fraction of cortex in the mouse brain (**Fig. 3**). Among its many substructures, agranular insular cortex (AI) is a preserved mammalian structure, while during evolution, especially in primates, the granular insular cortex (GI) expanded and exceeded the size of AI, while new subdivisions appear^62^. This is reflected in our observation that the fractional volume of AI in the insular cortex is 77% in mouse only 6% in the marmoset, together with the observation that the many insular cortical regions in the RIKEN atlas do not have corresponding regions in the Allen atlas. In the current analysis, we restricted the volume comparison to cerebral cortex because only the cerebral cortex was thoroughly segmented in both (Allen) mouse and (RIKEN) marmoset volumetric atlases. With future improvement of the whole-brain 3D reference atlases, similar analyses can be applied to other brain structures such as thalamus^59^ and brain stem^65^ quantitatively.

In recent years, studies have focused on addressing the challenge of cross-species whole-brain correspondence using quantitative methods. White matter tracts and myelin maps (T1w/T2w) in structural MRI^66,67^, as well as resting-state inter-regional correlation maps in functional MRI^68,69^ were used as vehicles for inferring organizations of brain structures. These are promising directions for establishing brain-wide homological relationships across species at the macroscopic level^70^, although caution needs to be taken in interpreting the signals^71^. Through the identification of a full set of common gray-matter leaf-level structures across atlases, individual brain regions are disentangled from disagreements arising from divergent hierarchical organizations imposed by experts, facilitating computational comparisons. Focusing on the leaf-level structures brings us closer to the spatially common coordinates employed across species, which proves advantageous both experimentally and computationally. Note that brain regions that were differently assigned into higher-level regions in the two atlases (**Extended data Table 1**) illustrate differences in the criteria used to establish the atlas hierarchies.

### Visualization tools for atlas hierarchy and homologous correspondences

The Brain Architecture Portal provides visualizations for tree hierarchies for the mouse brain atlas (http://brainarchitecture.org/mouse-connectivity-home), based on the Allen atlas, and the marmoset brain atlas (http://marmoset.brainarchitecture.org/), based on the RIKEN atlas. The brain region segmentation in the reference space is displayed in all three orthogonal views. By selecting a specific point in the reference space, the region information is displayed. With “Tree Search”, all brain regions from each atlas are displayed in a nested tree structure representing the respective hierarchy (**Fig. 4A**).

**Fig. 4.**
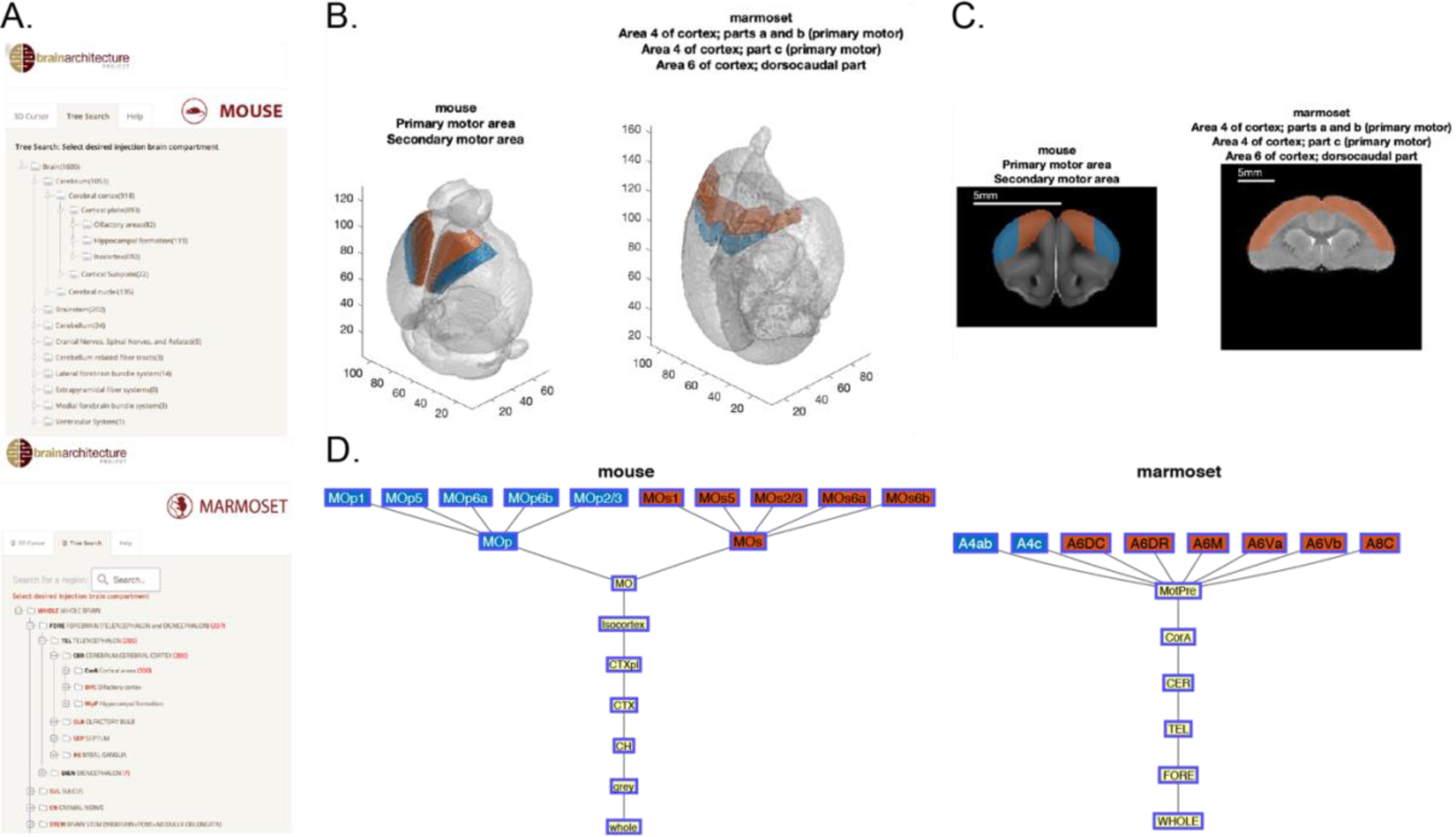
**A**. Hierarchical trees of Allen (top) and RIKEN (bottom) brain atlases taken from the Brain Architecture Portal (http://brainarchitecture.org). **B-D**. Sample outputs of the visualization tool for motor cortical areas. **B**. 3D views of the mouse reference brain (left, 100 µm isotropic) and marmoset reference brain (right, voxel size 240×240×224 µm^3^) with the primary and secondary motor areas labeled in blue and orange, respectively. Note that only 3 out of the 8 leaf-level structures in the marmoset brain are shown in the titles. **C**. Sample coronal sections through motor cortices (color labeled) in mouse (left) and marmoset (right). **D**. Primary (blue) and secondary (orange) motor cortex areas in the mouse (left) and marmoset (right) atlas hierarchies.

To enhance the accessibility of the framework developed in this study, and to provide an interface where results from this study can be easily queried, we developed a MATLAB-based query tool with both a command-line version and a GUI version (MATLAB App). In the command-line version, a series of prompts within a MATLAB interface helps guide the user through the process of specifying species and searching for a brain region (using the acronym, partial or full name) to be queried. If there is ambiguity, a list of candidate regions with similar names are presented to the user for precise identification of the brain region. To use the MATLAB App, the user selects from analogous dropdown lists the brain regions of either species to retrieve the corresponding homologous regions in the other species, with both volumetric and ontological tree outputs displayed. Note that both the App and command line tools assume the usage of acronyms or names corresponding to the Allen atlas and RIKEN volumetric atlases. As output, up to four figure panels appear, displaying 1) a dual view of both species’ reference brains in 3D, with queried region overlaid; and 2) a dual view of the coronal plane where the queried brain region is visible in both species’ reference brains; 3) the queried brain region’s hierarchy tree in the specified species’ atlas; and, 4) the corresponding brain region’s hierarchy tree in the counterpart species’ atlas. For visualization’s sake, references brains are not displayed to scale, but scale bars are presented as a part of the coronal plane images. When there is no correspondence found (Types O, U), no hierarchy tree or reference brain images in the other species are displayed. In the case where the queried brain region contains multiple leaf-level structures, all subregions are displayed, along with their cross-species correspondences. Colors of corresponding leaf-level regions are matched across the figures (**Fig. 4B-D**).

### Concluding remarks and future directions

Studying the brain with hierarchically organized atlases has advanced our understanding of neuroanatomy^35^. However, expert neuroanatomists disagree about how to group the smallest structures in the atlases, leading to different hierarchies being used within and between species. This makes comparative analyses challenging and inhibits cross-species data analysis platforms. We depart from the traditional hierarchical view and adopt the perspective that for homological analysis between different taxa, particularly concerning the mouse and marmoset model organisms in neuroscience, dealing with the most granular, leaf-level structures in the compartmental hierarchy is, in fact, more straightforward. We proceed with this approach and demonstrate how a working framework can be organized to compare mouse and marmoset brains, both at the level of compartment names, and also at the level of volumetric atlases. Our approach quantifies the degree and nature of correspondence between mouse and marmoset brain atlases and provides a concrete computational platform for future analyses.

Our framework for comparing leaf-level structural homology has various applications in modern comparative neuroanatomy. A key application is deriving comparable mesoscale connectivity across species, allowing for the comparison of neuron projections^72^ while preserving species-typical connectivity patterns^18^. By identifying corresponding brain structures in 3D reference spaces for both species, we can quantitatively characterize common and species-specific projection patterns. Additionally, our framework can be used as a central platform for analyzing gene expression data in mouse^44^ and marmoset brains^73^ for cell type composition of individual regions, together with histological data of cytoarchitecture and connectivity^74,75^. This comprehensive analysis can provide evidence of homologous brain regions and identify convergent and species-specific mesocircuit motifs. Furthermore, our method, exemplified by the correspondence between mouse and marmoset adult brain structures (Table 1), can be extended to developing brains and other species to expand the scope of comparative neuroanatomy.

## Methods

### Marmoset and mouse reference atlases

To maximize the utility of this framework, we adopted nomenclature and anatomical organization of brain regions in mouse and marmoset from the most commonly used atlases in the neuroanatomy field and publicly available in digital format. For the mouse reference atlases, we used the Allen Mouse Brain Atlas^44^ (http://download.alleninstitute.org/informatics-archive/), which followed the same concept as Swanson’s rodent brain organization^53^, and the Paxinos Mouse Atlas (**Extended data Table 2**, row 2). For the marmoset brain reference atlas, we used the Brain/MINDS atlas^45^ (https://dataportal.brainminds.jp/), which is almost entirely a subset of the Paxinos marmoset brain atlas^54^; this was used to obtain a complete set of leaf-level brain regions, which was not possible in the RIKEN atlas. The hierarchical structure of each atlas was visualized as a tree using GraphViz (AT&T) (**Figures 1, 2**). The root is the whole brain, and the leaves are the granular level of brain regions. In all plotted hierarchical trees, only gray matter was shown.

Volumetric atlases used were the Allen Mouse Brain Atlas^43^ and the RIKEN marmoset atlas. The RIKEN marmoset atlas only annotates ∼60% of all gray matter at a level beyond parent region designations (e.g. cerebellum). Therefore, we merged the RIKEN marmoset atlas with two additional marmoset brain volumes with partial segmentation: the thalamic, hippocampal, midbrain, brainstem, and cerebellar atlas from Saleem et al. (**Extended data Table 2**; row 6) and the NIH-Silva marmoset brain atlas for cortical refinements (**Extended data Table 2**, row 5). This allowed for close to complete segmentation, at a leaf-node level, of the marmoset brain. However, segmented brain volumes for both mouse and marmoset do not represent the full set of leaf-level nodes present in the nomenclatures, and so compartment name and volume based analyses, while similar in underlying philosophy, are presented separately in this manuscript.

### Brain region correspondence across species

Leaf-level structures were initially defined as brain compartments that cannot be further divided into substructures. The Allen atlas contains 1327 structures of the mouse brain, in which 1038 are candidates for leaf-level structures, the remainder pertaining to higher level groupings. The RIKEN atlas contains 800 structures of the marmoset brain, in which 689 are candidates for leaf-level structures. These were supplemented by the Paxinos mouse and Paxinos marmoset atlases. To identify corresponding leaf-level regions, we needed to 1) decide the appropriate “leaf level”, and 2) find homologically corresponding regions in the other species. Several criteria, in addition to no subregions in the atlas, were applied to decide leaf-level structures. Cortical layers were removed and collapsed into the corresponding cortical area. Homological correspondence was established while accounting for the notion that homologous structures may have lower-level elements that are not homologous^31^. All revisions to the leaf-level structures were carefully noted and verified manually via careful examination of the stack of 2D brain region annotations from the print atlases overlaid on tissue section images for each atlas.

To find correspondences, we first performed an initial “guess” based on the nomenclature itself by algorithmically searching for similarities in text strings. Highly similar terms such as “triangular septal nucleus” in the RIKEN marmoset atlas and “triangular nucleus of septum” in the Allen mouse atlas were taken as tentative correspondences. Next, we leveraged online databases such as NeuroNames (braininfo.org) to assist the literature-based search. All correspondences were verified or manually curated by neuroanatomy experts in the team, to bring in additional literature and professional judgement. In addition to the criteria as described above, some corresponding areas were topographically mapped based on the direct comparison of the atlases, together with any information extracted from the rat atlas of Swanson and a human atlas^76^, whenever possible, plus other supporting references found in the literature.

Final leaf-level structures for analysis satisfied the conditions that: 1) no correspondence could be found in any substructures, and 2) they are the most granular level structures for which correspondences could be found. The complete list of correspondences with the references are presented in **Supplementary Table 1**. The list of final leaf-level structures only interpolated across both Allen and Paxinos mouse atlases for the mouse ontology can be found in **Supplementary Tables 2 & 3**.

### Transplanting hierarchies on key regions

“Key regions” were defined as leaf-level structures that were common to both Allen mouse and RIKEN marmoset atlases, in other words, regions with Type I – IV correspondences. To demonstrate the Allen mouse atlas hierarchy, the key regions were identified from the Allen atlas as leaf-level structures. The relational map between leaf-level structures and higher-order structures from the Allen atlas were then applied to these key regions. Note that leaf-level structures in the Allen atlas that did not have corresponding regions in the marmoset brain (Types O, U) were not included in this process. Conversely, the RIKEN atlas hierarchy was applied to the key regions, with higher-order structures from the RIKEN atlas. Similarly, leaf-level structures in the RIKEN marmoset atlas that did not find correspondence in the mouse brain were excluded.

### Brain volume comparison

Volumetric analyses of brain region compositions and comparisons between corresponding structures are based on 3D reference atlases of mouse and marmoset. The Allen Mouse Brain Atlas provided a 3D segmentation of the entire brain, or Common Coordinate Framework version 3 (CCF3)^56^ in 25 µm isotropic volume. The marmoset reference brain came from an average of 43 female marmosets via *in vivo* high resolution T2 mapping, with 110 µm isotropic volume, mapped to a common reference space. Brain region segmentation was refined based on existing digital atlases^57,77,78^ to cover the entire brain. For quantitative volumetric analyses we merged the RIKEN marmoset atlas with an additional marmoset brain volume with partial segmentation, the thalamic, hippocampal, midbrain, brainstem, and cerebellar atlas from Saleem et al. (**Extended data Table 2**, row 6). This allowed for close to complete segmentation, at a leaf-node level, of the marmoset brain. Briefly, the 3D reference volumes of the Brain/MINDS marmoset reference brain and the NIH and Saleem marmoset reference brains were co-registered, mapped to the reference space, and the segmentations were combined. The refined marmoset brain atlas achieved >87% leaf-level segmentation of the reference volume, excluding white matter and ventricles. Of note, leaf-level regions again had to be selected for volumetric comparison, as labeled regions comprise only a subset of those present in mouse and marmoset nomenclatures; we obtained 337 mouse and 319 marmoset leaf-level regions in the labeled volumes we utilized. Volumetric comparisons were established between gray matter labels for each species. Percentages of all labeled gray matter volume, including general labels, were calculated for each relation type. We additionally calculated the fraction of volumetrically labeled leaf-level region names per relation type.

### Volumetric merging of marmoset brain atlas segmentations

The algorithmic step of merging marmoset brain atlas segmentations, using the RIKEN marmoset atlas as the primary segmentation, proceeded as a 5-step process. First, NIH-Silva and Saleem et al. reference space volumes were registered with, and mapped into, our marmoset reference space based on the averaging of 43 in vivo volumetric MRIs. Second, the transforms necessary to take NIH-Silva and Saleem et al. reference volumes into the same space as the RIKEN marmoset segmentation were applied to each of these segmented volumes. Third, we calculated overlap between region IDs in each segmented volume for each atlas. Because the NIH-Silva atlas provided refinements in cortical areas and the Saleem et al. atlas provided refinements in subcortical, midbrain and hindbrain regions, we treated these two atlases as purely refinements on the RIKEN marmoset atlas, and disjoint sets from one another. The overlaps were therefore only calculated between each of the additional two segmented volumes and the RIKEN marmoset atlas volume. We treat these overlaps as analogous to conditional probabilities that a region from one segmentation maps to a given RIKEN marmoset segmentation. Fourth, we took the maximum conditional probability of mapping to a RIKEN marmoset segmentation from each of the NIH-Silva and Saleem et al. segmentations and generated concordance matrices between each of these segmentations and the RIKEN volume. Fifth, this allowed us to generate a bipartite graph of these concordances, which was then subsetted to only cortical segmentations providing refinement relative to the RIKEN parcellations for the NIH-Silva atlas, and only hippocampal, thalamic, midbrain, cerebellar and brainstem segmentations providing refinement relative to RIKEN parcellations for the Saleem et al. atlas. The above procedure accomplished algorithmic refinement of high-level “cerebrotypes”^79^ including telencephalon, diencephalon, midbrain, medulla, pons, cerebellum and circumventricular organs. The resultant combined volume was then manually proofread for anatomical correctness and gap filling in the grey matter.

### Query tool

The MATLAB-based visualization tool was developed using MATLAB R2022a (MathWorks) on macOS version 14. A command-line version and a MATLAB App are released with this manuscript, also available on GitHub (https://github.com/bingxinghuo/brainatlascomparison).

## Supporting information

Supplementary Table 1: An excel file listing all correspondences between all named mouse and marmoset regions

Supplementary Table 2: An excel file with correspondences from mouse to marmoset, listed as a unified table prepared for analytics

Supplementary Table 3: An excel file with correspondences from marmoset to mouse, listed as a unified table prepared for analytics

Supplementary Tables 4 & 5, plus references for Supplementary Table 1

## Data availability

Data is available in excel and csv tables provided as linked documents in the Supplementary Information section. Supplementary Information datasets/tables are as follows:

1. **Supplementary Table 1**: An excel file listing all correspondences between all named mouse and marmoset regions, collated across Allen Mouse, Paxinos Mouse, Paxinos Marmoset, RIKEN Marmoset, NIH-Silva Marmoset, and Saleem et al. Marmoset atlases, whose references are collated together in **Extended Data Table 2**. References for making anatomical inferences given the atlas nomenclatures can also be found in **Extended Data** under the **Supplementary Table 1 References** section.
2. **Supplementary Table 2**: An excel file with correspondences from mouse to marmoset, listed as a unified table prepared for analytics. This was used to generate **Tables 2 & 3**.
3. **Supplementary Table 3**: An excel file with correspondences from marmoset to mouse, listed as a unified table prepared for analytics. This was used to generate **Tables 2 & 3**.

## Code availability

The MATLAB App for volumetric and ontological comparisons, both the package and the code, are available for download via GitHub: https://github.com/bingxinghuo/DAP/tree/master/AtlasComparison/cross-species_atlas

## Acknowledgements.

The authors would like to thank Drs. Angela Roberts, Tadashi Isa, Marcello Rosa, and Katherine Rockland on helpful discussions on the cross-species homological correspondences. We also acknowledge and are grateful for the generous support of the National Institutes of Health towards this project, on awards number **UG3MH126869-01** and **UM1NS132173**. We also acknowledge the generous support of the Crick-Clay Professorship, Cold Spring Harbor Laboratory.

## Author contributions

***Bingxing Huo*** helped initiate the study and analytic pipelines, designed and implemented the Matlab app, contributed to the major comparative table, **Supplementary Table 1**, helped compile the references for the manuscript and **Supplementary Table 1**, designed and drafted **Figures 1-4**, and was one of the writers of the manuscript.

***Christopher Mezias*** helped design the analytics pipelines, implemented the analytics pipelines and calculated results, contributed to the major comparative table, **Supplementary Table 1**, designed and drafted **Tables 1-3**, and was one of the writers of the manuscript.

***Mihail Bota*** was the primary contributor towards **Supplementary Table 1**, helped compile the references for the manuscript and **Supplementary Table 1** via literature search, and helped revise the manuscript text.

***Jaikishan Jayakumar*** contributed to and helped edit the major comparative artifact, **Supplementary Table 1** and helped revise manuscript text and figure and table layouts.

***Partha Mitra*** proposed the study and analytics pipeline design, helped revise the major comparative artifact associated with the manuscript, **Supplementary Table 1**, and wrote and revised manuscript text. He oversaw and directed the overall project.

## Materials & Correspondence

All requests for materials and inquiries on the manuscript should be addressed to. Code and datasets are available as indicated in the **Data Availability** and **Code Availability** sections.

