## Supplementary Tables 4 & 5, plus references for Supplementary Table 1 for "Establishing neuroanatomical correspondences across mouse and marmoset brain structures"

### Extended Data Figures & Tables

**Extended data Table 1**. Leaf-level structures are assigned to high-level regions differently in the two atlases. Abbreviations: TEL: telencephalon/cerebrum; DIEN: diencephalon/interbrain; MID: midbrain; PONS: pons; MED: medulla oblongata; CvO: circumventricular organs.

| | Marmoset | | | Mouse | | |
|---|---|---|---|---|---|---|
| **Type** | **Acronym** | **Region name** | **Cerebrotype** | **Acronym** | **Region name** | **Cerebrotype** |
| 1 | SNC | Substantia nigra; compact part | **TEL** | SNc | Substantia nigra, compact part | **MID** |
| 1 | SNL | Substantia nigra; lateral part | **TEL** | SNl | Substantia nigra, lateral part | **MID** |
| 1 | SNR | Substantia nigra; reticular part | **TEL** | SNr | Substantia nigra, reticular part | **MID** |
| 1 | STh | Subthalamic nucleus | **TEL** | STN | Subthalamic nucleus | **DIEN** |
| 1 | E | Ependyma & subependymal layer | **TEL** | SEZ | subependymal zone | **CvO** |
| 1 | APT | Anterior pretectal nucleus | **DIEN** | APN | Anterior pretectal nucleus | **MID** |
| 1 | MPT | Medial pretectal area | **DIEN** | MPT | Medial pretectal area | **MID** |
| 1 | OT | Nucleus of the optic tract | **DIEN** | NOT | Nucleus of the optic tract | **MID** |
| 1 | OPT | Olivary pretectal nucleus | **DIEN** | OP | Olivary pretectal nucleus | **MID** |
| 1 | PrC | Precommissural nucleus | **DIEN** | PRC | Precommissural nucleus | **MID** |
| 2 | MCPC,PCom | Magnocellular nucleus of the posterior commissure,Nucleus of the posterior commissure | **DIEN, MID** | NPC | Nucleus of the posterior commissure | **MID** |
| 1 | SCO | Subcommissural organ | **CvO** | SCO | Subcommissural organ | **MID** |
| 1 | PP | Peripeduncular nucleus | **MID** | PP | Peripeduncular nucleus | **DIEN** |
| 1 | ME | Median eminence | **CvO** | ME | Median eminence | **DIEN** |
| 1 | Pi | Pineal gland | **CvO** | PIN | Pineal body | **DIEN** |
| 1 | SFO | Subfornical organ | **CvO** | SFO | Subfornical organ | **DIEN** |
| 1 | VOLT | Vascular organ of the lamina terminalis | **CvO** | OV | Vascular organ of the lamina terminalis | **DIEN** |
| 1 | ATg | Anterior tegmental nucleus | **PONS** | AT | Anterior tegmental nucleus | **MID** |
| 1 | 6N | Abducens nucleus | **PONS** | VI | Abducens nucleus | **MED** |
| 1 | DC | Dorsal cochlear nucleus | **PONS** | DCO | Dorsal cochlear nucleus | **MED** |
| 1 | GrC | Granule cell layer of the cochlear nuclei | **PONS** | CNlam | Granular lamina of the cochlear nuclei | **MED** |
| 2 | VCA,VCP | Ventral cochlear nucleus; anterior part,Ventral cochlear nucleus; posterior part | **PONS** | VCO | Ventral cochlear nucleus | **MED** |
| 2 | 7N,7SH | Facial nucleus,Facial motor nucleus; stylohyoid part | **PONS** | VII | Facial motor nucleus | **MED** |
| 1 | Tz | Nucleus of the trapezoid body | **PONS** | NTB | Nucleus of the trapezoid body | **MED** |
| 1 | AP | Area postrema | **CvO** | AP | Area postrema | **MED** |
| 1 | PBG | Parabigeminal nucleus | **PONS** | PBG | Parabigeminal nucleus | **MID** |
| 1 | PTg | Pedunculotegmental nucleus | **PONS** | PPN | Pedunculopontine nucleus | **MID** |
| 1 | CLi | Caudal linear nucleus of the raphe | **PONS** | CLI | Central linear nucleus raphe | **MID** |
| 1 | RMg | Raphe magnus nucleus | **PONS** | RM | Nucleus raphe magnus | **MED** |
| 1 | Sag | Sagulum nucleus | **PONS** | SAG | Nucleus sagulum | **MID** |
| 1 | LT | Lateral terminal nucleus of the accessory optic tract | **MED** | LT | Lateral terminal nucleus of the accessory optic tract | **MID** |
| 1 | KF | Kolliker-Fuse nucleus | **MED** | KF | Koelliker-Fuse subnucleus | **PONS** |
| 1 | SuS | Superior salivatory nucleus | **MED** | SSN | Superior salivatory nucleus | **PONS** |

**Extended data Table 2**. A list of the 5 core atlas resources used for ontological, volumetric, or both sets of mouse-marmoset anatomical comparisons in the manuscript.

| **Atlas Abbr. Name** | **Atlas Citation** |
|---|---|
| Allen Mouse Atlas | Wang, Q. *et al.* The Allen Mouse Brain Common Coordinate Framework: A 3D Reference Atlas. *Cell* **181**, 936-953.e20 (2020). |
| Paxinos Mouse Atlas | Paxinos, G. & Franklin, K. The Mouse Brain in Stereotaxic Coordinates. (2004). |
| Paxinos Marmoset Atlas | Paxinos, G., Watson, C., Petrides, M., Rosa, M. & Tokuno, H. *The Marmoset Brain in Stereotaxic Coordinates*. (Academic Press, 2012). doi:10.1016/j.brainresbull.2013.01.009. |
| RIKEN Marmoset | Hashikawa, T. *et al.* The 3-Dimensional Atlas of the Marmoset Brain, Reconstructible in Stereotaxic Coordinates. *Brain Sci* 33–329 (2018) doi:10.1007/978-4-431-56612-0_2. |
| NIH-Silva Marmoset | Liu, C. *et al.* A digital 3D atlas of the marmoset brain based on multi-modal MRI. *NeuroImage* **169**, 106–116 (2018). |
| Saleem Marmoset | Saleem KS, Avram AV, Yen CC, Magdoom KN, Schram V, Basser PJ. Multimodal anatomical mapping of subcortical regions in Marmoset monkeys using high-resolution MRI and matched histology with multiple stains. bioRxiv [Preprint]. 2023. doi: 10.1101/2023.03.30.534950. PMID: 37034636; PMCID: PMC10081239. |

**Supplementary Table 1 References**

21. Jones EG. 2007. The Thalamus 2 Volume Set, Cambridge University Press. Cambridge University Press.

22. Jones EG, Rubenstein JLR. 2004. Expression of regulatory genes during differentiation of thalamic nuclei in mouse and monkey. *J Comp Neurol* **477**:55–80. doi:10.1002/cne.20234

23. Kaas JH. 2010. The Auditory Cortex 407–427. doi:10.1007/978-1-4419-0074-6_19

24. Krubitzer LA, Kaas JH. 1990. The organization and connections of somatosensory cortex in marmosets. *The Journal of neuroscience : the official journal of the Society for Neuroscience* **10**:952–974.

25. Lein ES, Hawrylycz MJ, Ao N, Ayres M, Bensinger A, Bernard A, Boe AF, Boguski MS, Brockway KS, Byrnes EJ, Chen Lin, Chen Li, Chen T-M, Chin MC, Chong J, Crook BE, Czaplinska A, Dang CN, Datta S, Dee NR, Desaki AL, Desta T, Diep E, Dolbeare TA, Donelan MJ, Dong H-W, Dougherty JG, Duncan BJ, Ebbert AJ, Eichele G, Estin LK, Faber C, Facer BA, Fields R, Fischer SR, Fliss TP, Frensley C, Gates SN, Glattfelder KJ, Halverson KR, Hart MR, Hohmann JG, Howell MP, Jeung DP, Johnson RA, Karr PT, Kawal R, Kidney JM, Knapik RH, Kuan CL, Lake JH, Laramee AR, Larsen KD, Lau C, Lemon TA, Liang AJ, Liu Y, Luong LT, Michaels J, Morgan JJ, Morgan RJ, Mortrud MT, Mosqueda NF, Ng LL, Ng R, Orta GJ, Overly CC, Pak TH, Parry SE, Pathak SD, Pearson OC, Puchalski RB, Riley ZL, Rockett HR, Rowland SA, Royall JJ, Ruiz MJ, Sarno NR, Schaffnit K, Shapovalova NV, Sivisay T, Slaughterbeck CR, Smith SC, Smith KA, Smith BI, Sodt AJ, Stewart NN, Stumpf K-R, Sunkin SM, Sutram M, Tam A, Teemer CD, Thaller C, Thompson CL, Varnam LR, Visel A, Whitlock RM, Wohnoutka PE, Wolkey CK, Wong VY, Wood M, Yaylaoglu MB, Young RC, Youngstrom BL, Yuan XF, Zhang B, Zwingman TA, Jones AR. 2007. Genome-wide atlas of gene expression in the adult mouse brain. *Nature* **445**:168–176. doi:10.1038/nature05453

50. Smith JB, Alloway KD, Hof PR, Orman R, Reser DH, Watakabe A, Watson GDR. 2018. The relationship between the claustrum and endopiriform nucleus: A perspective towards

consensus on cross-species homology. *J Comp Neurol* **527**:476–499. doi:10.1002/cne.24537
